# Reactivation of an Embryonic Cardiac Neural Crest Transcriptional Subcircuit During Zebrafish Heart Regeneration

**DOI:** 10.1101/2025.01.16.633462

**Authors:** Rekha M. Dhillon-Richardson, Alexandra K. Haugan, Luke W. Lyons, Joseph K. McKenna, Marianne E. Bronner, Megan L. Martik

**Affiliations:** Department of Molecular and Cell Biology, University of California, Berkeley; Berkeley, CA, USA; Division of Biology, California Institute of Technology; Pasadena, CA, USA

**Author notes:** Authors contributed equally (Rekha M. Dhillon-Richardson, Alexandra K. Haugan).

## Abstract

During vertebrate development, the heart primarily arises from mesoderm, with crucial contributions from cardiac neural crest cells that migrate to the heart and form a variety of cardiovascular derivatives. Here, by integrating bulk and single cell RNA-seq with ATAC-seq, we identify a gene regulatory subcircuit specific to migratory cardiac crest cells composed of key transcription factors *egr1, sox9a, tfap2a* and *ets1*. Notably, we show that cells expressing the canonical neural crest gene *sox10* are essential for proper cardiac regeneration in adult zebrafish. Furthermore, expression of all transcription factors from the migratory cardiac crest gene subcircuit are reactivated after injury at the wound edge. Together, our results uncover a developmental gene regulatory network that is important for cardiac neural crest fate determination, with key factors reactivated during regeneration.

**SIGNIFICANCE:** Many common human congenital heart defects are linked to problems that arise during cardiac neural crest development. Here, we use the zebrafish, which has the remarkable ability to regenerate their adult heart, to understand the genetic programs that control cardiac development and adult repair. We discover a set of genes that control the development of the cardiac neural crest and find that these genes are reactivated after heart injury in the adult zebrafish. Unlike the zebrafish, human hearts have a very limited ability to regenerate after injury. Our findings in zebrafish can provide insight to potential clinical interventions for congenital heart defects and adult heart damage.

## INTRODUCTION

The neural crest (NC) is a multipotent, highly migratory cell population that arises early in vertebrate embryonic development. Distinct NC populations exist along the body axis and vary in their developmental potential to give rise to certain neural crest derivatives^1–3^. Among these unique axially-restricted populations is the cardiac neural crest (CdNC), which is essential for proper cardiovascular system development. The CdNC contributes to several cardiac cell types and structures, including cardiomyocytes, coronary vessels, the interventricular septum, the outflow tract cushions, pharyngeal arch arteries, valves, cardiac fibroblasts, and cardiac ganglia^4–10^.

In the developing zebrafish embryo, premigratory CdNC cells reside in the dorsal neural tube from rhombomere 1 - somite 6 by 17 hours post fertilization (hpf)^7^. Between 17-24hpf, the CdNC migrates to the circumpharyngeal area of forming pharyngeal arches 1 and 2^11^. These cells then continue their migration into the developing myocardium, ultimately giving rise to ∼12-13% of total cardiomyocytes in zebrafish^7,11–13^. Lineage tracing experiments in the mouse and chick have shown that the CdNC contribution to cardiomyocytes is similar in amniotes^2^.

Disruption of CdNC formation has substantial effects on heart development. Ablation of premigratory CdNC in zebrafish causes defects in heart looping and reduces ventricular function at 48 hpf^12^. Genetic ablation of the zebrafish NC results in a reduced number of total cardiomyocytes, defective myocardial maturation, and abnormal atrioventricular valve development^11^. Furthermore, specific ablation of embryonic NC-derived cardiomyocytes in zebrafish results in severe heart defects in the adult, including hypertrophic cardiomyopathy and heart failure during stress tests^14^. Thus, the CdNC is essential for proper cardiac development and function in the zebrafish.

In amniotes, ablation studies show that loss of the CdNC leads to a wide range of cardiac defects, including persistent truncus arteriosus, abnormal myocardium function, and misalignment of the arch arteries^4,15^. These defects are highly reminiscent of some of the most common human congenital heart defects. Therefore, understanding CdNC development in diverse animal systems has implications for human health. The formation of distinct axial populations of the NC, like the CdNC, are regulated by unique gene regulatory networks (GRNs). For example, in the chicken embryo, a gene subcircuit comprised of transcription factors *Tgif1, Sox8*, and *Ets1* was shown to regulate CdNC axial specification. When CdNC cells are ablated in the chick, trunk neural crest (TNC) cells reprogrammed with this subcircuit can be grafted in place of the CdNC and rescue ablation defects.^16^. Thus, the potential use of axial-specific NC gene networks holds promise for reprogramming and repair and remains an intriguing avenue for therapeutic interventions.

The critical importance of the embryonic CdNC for proper heart formation raises the intriguing possibility that CdNC-derived cells may also play a role in adult heart repair. Consistent with this hypothesis, key NC markers like *sox10* and *tfap2a* are upregulated in adult regenerating zebrafish cardiac tissue as shown by *in situ* hybridization and RNA sequencing^2^. Additionally, *sox10*-expressing cardiomyocytes in the adult zebrafish preferentially proliferate during heart regeneration relative to other cardiomyocytes^2,17^. In mice, which have the ability to regenerate their hearts for the first week of life, myelin protein-0 (P0) Cre lineage tracing has shown that putative NC stem cells gather at the ischemic border zone after myocardial infarction, indicating a potential role for these cells in regeneration^18,19^. However, further investigation is needed as P0 does not exclusively label NC stem cells. Overall, these data suggest that CdNC-derived cells, and potentially CdNC genes, could have a unique role in cardiac regeneration.

To better understand the formation of the migratory CdNC in zebrafish, we have assembled a gene regulatory subcircuit that underlies its development. We profile the CdNC to uncover a core set of transcription factors and cis-regulatory elements active specifically in this population. Importantly, enhancer analyses and CRISPR knockouts reveal a novel role for transcription factor *Egr1* in promoting the expression of key CdNC transcription factors, highlighting its significance in CdNC gene regulation. Additionally, we explore the role these CdNC markers may play in adult cardiac regeneration, identifying a set of migratory CdNC transcription factors that are reactivated in and around the regenerating tissue after injury in adult zebrafish.

## RESULTS

### Uncovering the distinct gene expression signature of zebrafish cardiac neural crest cells

To explore the unique transcriptional profile of migratory CdNC cells, we performed bulk RNA sequencing (RNA-seq) on CdNC, TNC and non-NC populations for differential expression analysis. To label the migratory CdNC, we used a transgenic line using the sox10 promoter driving GAL4/UAS-mediated expression of *mCherry* (*Tg(−4.9sox10:GAL4-UAS-cre;UAS:NfsB-mCherry;myl7:nucGFP)*, herein referred to as *Tg(sox10:Nfsb-mCherry)*)^11^. Specifically, an enriched population of migratory CdNC was isolated by dissecting the midbrain-hindbrain boundary to somite 6 of 16 somite-stage (ss) embryos followed by fluorescence-activated cell sorting (FACS) to capture *sox10*-expressing cells^7^ (Fig. S1A). Non-NC cells from the same axial level were also collected via FACS as a negative population for differential expression analysis. CdNC RNA-seq libraries were then also compared to a TNC dataset produced by FACS on dissected tissue of 24 hpf embryos from somite 7 to somite 16 of *Tg*(*-4.9sox10:eGFP)*^20^ (Fig. S1A). 591 genes were upregulated in the CdNC compared to non-NC tissue, and 2573 genes were unique to the CdNC when compared to the migratory TNC (log2 fold change >1; p-adj<0.05) (Table S1). As expected, canonical NC genes such as *sox10, tfap2a*, and *ets1* were upregulated in the NC (*sox10+*) population compared to non-NC (*sox10-*) tissue, reaffirming that we were enriching for CdNC with our collection scheme (Fig. S2A). Both transcription factors and signaling molecules were upregulated in the CdNC compared to the TNC, including previously identified NC genes *mafba, prdm1a, egr1, foxc1b, sox9a, fli1*, and *cxcr4b*^16,21–26^ (Fig. S2B).

To further resolve the heterogeneity of regulatory states within the developing CdNC, we performed single cell RNA sequencing (scRNA-seq), utilizing the same scheme as bulk RNA-seq for isolating enriched populations of CdNC (Fig. S1A). We prepared and sequenced 3,828 cells at an average depth of 80,000 reads per cell using the 10X Genomics pipeline. Cell Ranger was used to exclude low quality cells and map reads to GRCz11. Scanpy, a Python-based computational tool for analyzing scRNA-seq data, was used to perform dimensionality reduction, Leiden clustering, and downstream analysis^27^.

We identified several broad cell type clusters within the CdNC population, including 2 clusters of developing pigment cells, 2 neuronal-like clusters, 1 ectomesenchymal cranial NC cluster, and 5 additional migratory NC clusters. Pigment cells were identified by expression of *mitfa* and *tyr* and neuronal cell types with *elavl3* and *gfap* (Fig. 1A, S1B). Ectomesenchymal cranial NC was recognized via the upregulation of ectomesenchymal genes (*prrx1a* and *twist1a/b*) and the expression of collagen-encoding genes (*col5a1, col1a2, col2a1b*) (Fig. 1A, S1B). The results revealed 12 clusters in total, suggesting that the migratory NC population was already quite heterogeneous in cell identity at 16ss.

**Figure 1.**
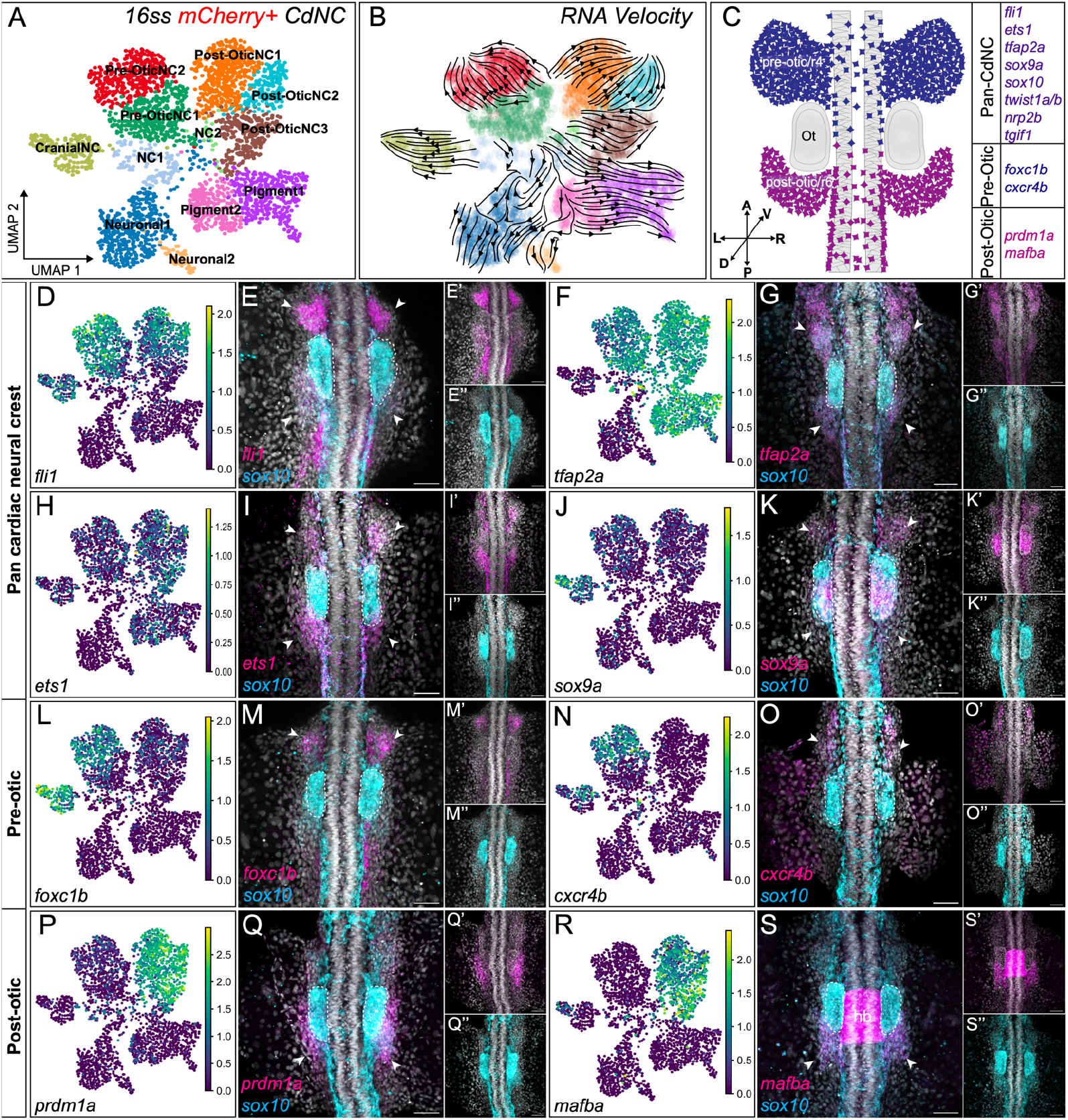
Single-cell analysis reveals two transcriptionally distinct populations of CdNC. **(A)** Labeled leiden clustering UMAP of scRNA sequencing of FAC sorted mCherry+ CdNC cells. **(B)** RNA velocity embedded on UMAP. **(C)** Illustration of a dorsal view, 16ss zebrafish embryo with pre and post otic CdNC domains labeled and validated genes listed that are restricted to either domain or shared amongst both domains. **(D-S’’)** UMAPs for individual genes and corresponding 20x confocal images of HCR expression patterns of dorsal view, 16ss zebrafish embryos, showing overlap in expression and individual channel insets of *sox10* in cyan and the following genes of interest in magenta: **(D-E’’)** *fli1* (n=11), **(F-G’’)** *tfap2a* (n=8), **(H-I’’)** *ets1* (n=8), **(J-K’’)** *sox9a* (n=6), **(L-M’’)** *foxc1b* (n=3), **(N-O’’)** *cxcr4b* (n=8), (P-Q’’) *prdm1a* (n=6), and (R-S’’) *mafba* (n=15). The white arrows indicate expression in pre-otic and/or post-otic CdNC streams, and the white dotted circles outline the otic vesicle. Acronyms: NC, neural crest; r4, rhombemere 4; r6, rhombemere 6; Ot, otic vesicle. Position: A, anterior; P, posterior; D, dorsal; V, ventral; L, left; R, right; hb, hindbrain. All scale bars are 50 um.

We used scVelo, a tool that leverages the ratio of spliced to unspliced mRNA transcripts to compute the RNA velocity for each gene, to assess cell state dynamics^28^. These velocities were then combined to estimate the future state of a given cell and projected onto the UMAP (Fig. 1B). Knowing that the migratory CdNC is a temporally heterogeneous cell population in which cells delaminate and begin their migration in waves, this analysis further confirmed that we captured a diverse population consisting of multiple cell states at a single time point. This also enabled us to refine cluster identity, as many clusters represented the same cell types in different regulatory states. From this, we inferred that the 5 additional migratory NC clusters represented temporally, spatially, and transcriptionally distinct groups of NC (Fig. 1A).

To validate cluster identity, we performed multiplexed hybridization chain reaction (HCR) for top markers in the 5 clusters^29^ (Fig. 1C). We found that *cxcr4b* and *foxc1b* were expressed in the rhombomere 4/pre-otic migratory stream, while *prdm1a* and *mafba* were predominantly expressed in the rhombomere 6/post-otic stream (Fig. 1L-O’’, 1P-S’’). Several genes, including *ets1, sox9a, tfap2a, fli1, tgif1, nrp2b, twist1a*, and *twist1b*, were expressed in both pre and post-otic streams of the CdNC (Fig. 1D-K’’, S2C-J’’). Thus, our transcriptomic datasets allowed us to identify unique molecular signatures of pre- and post-otic migratory CdNC.

### Identification of putative enhancer regions and construction of a migratory CdNC subcircuit

To investigate the regulation of these validated CdNC genes, we next profiled open chromatin using assay for transposase-accessible chromatin followed by sequencing (ATAC-seq). Utilizing the same scheme to isolate enriched populations of NC cells as the RNA sequencing experiments, we collected 16ss CdNC (mCh+) and non-NC (mCh-) cells (Fig. S1A). To uncover CdNC-enriched regions of open chromatin, we compared differential accessibility between CdNC and non-NC tissue using Diffbind^30,31^ (Fig. 2A). This analysis revealed 1,247 regions of open chromatin that were more accessible in the migratory CdNC as compared to non-NC cells (FDR< 0.05, Log2 Fold Change > +1) (Table S2). To identify the top transcription factor binding motifs within our CdNC-specific accessible chromatin, we performed *de novo* motif analysis using HOMER, which revealed enrichment of predicted binding sites for several CdNC transcription factors, including Tfap2a, Sox10, Ets1, Egr1, and Sox9^32^ (Fig. 2B). These data further support the conclusion that these transcription factors play a pivotal role in regulating CdNC identity. Aiming to understand the regulation of these core transcription factors, we examined peaks in proximity to each of these genes as putative enhancers in the CdNC-enriched regions. This revealed multiple differentially accessible regions near the *tfap2a, ets1, sox9a*, and *sox10* loci (Fig. 2C). Notably, all CdNC-enriched peaks near the *sox10* locus were previously identified and validated as NC enhancers in zebrafish, with peak 5 (here referred to as -22kb_E3) being critical for maintaining embryonic *sox10* expression^33^.

**Figure 2.**
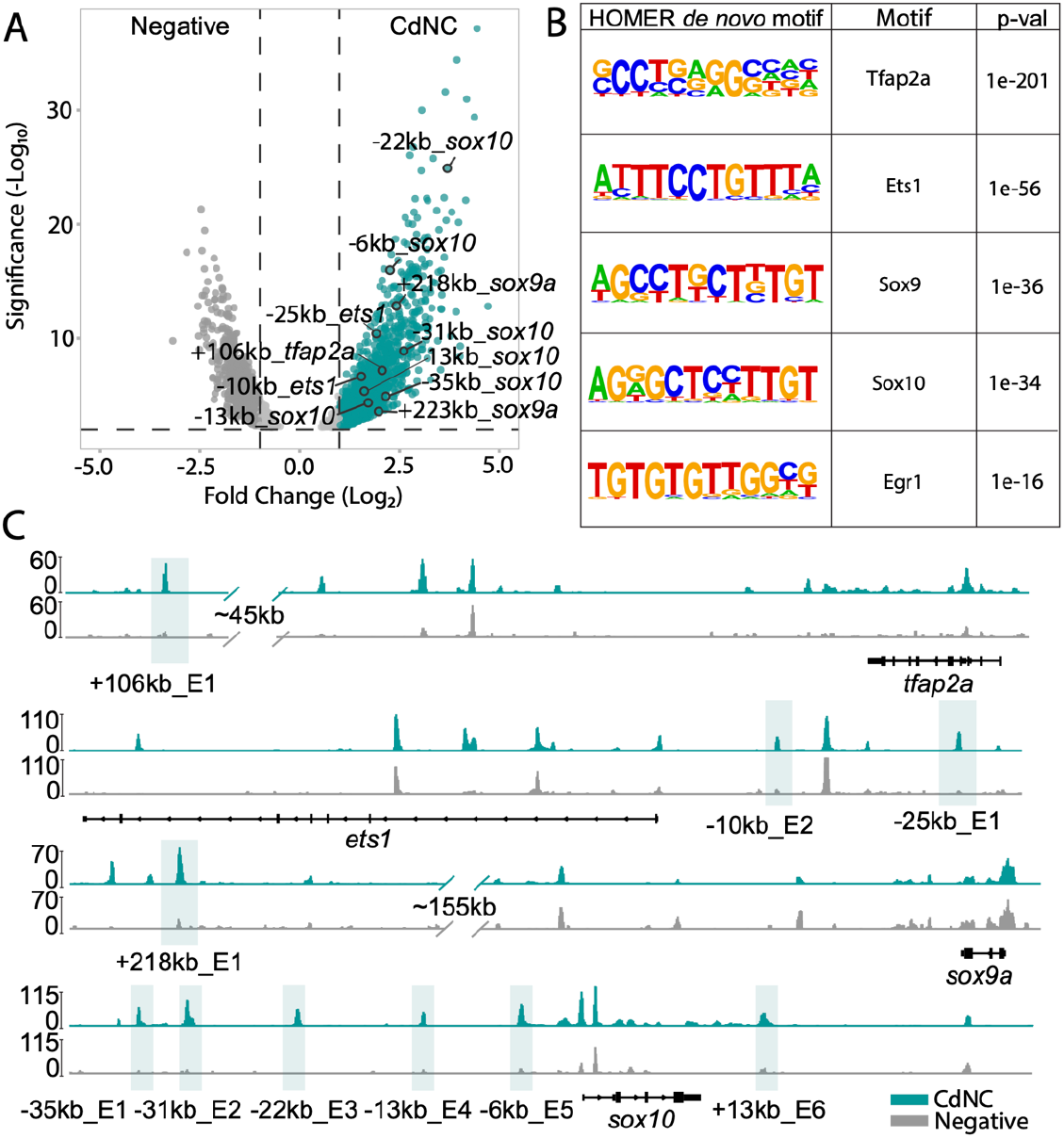
ATAC-seq and differential chromatin accessibility and transcription factor motif analysis uncovers putative enhancers and transcriptional regulators unique to the CdNC. **(A)** Volcano plot of merged peaks of sorted bulkATAC-seq shows differentially accessible regions in CdNC (mCherry+) and non-NC/negative cells (mCherry-). **(B)** Results of HOMER *de novo* motif analysis showing motif, transcription factor, and associated p-value. **(C)** IGV genome browser tracks of the *ets1, tfap2a, sox9a*, and *sox10* loci. Highlighted peaks represent putative enhancer regions called by Diffbind (turquoise track= CdNC, gray track = negative).

To identify probable transcription factor binding sites, these putative enhancer sequences were scanned using Find Individual Motif Occurrences (FIMO), revealing predicted binding sites for numerous NC transcription factors (Tfap2a, Jun, Twist1, Foxd3, Tgif1, Foxc1, Prdm1, and Mafb)^34^ (SFig. 3, Table S3). Many of these regions also contained predicted binding sites for Egr1 (SFig. 3). This was particularly interesting as the role of Egr1 in NC biology is not well characterized. Previous *in situ* hybridization experiments showed that *Egr1* was expressed in the migratory CdNC in chick, and, in zebrafish, *egr1* expression in the endoderm facilitates BMP signaling in cranial NC cells^16,35^. Furthermore, *egr1* is upregulated in migratory CdNC compared to TNC in our bulk RNA-seq data (Fig. S2B).

**Figure 3:**
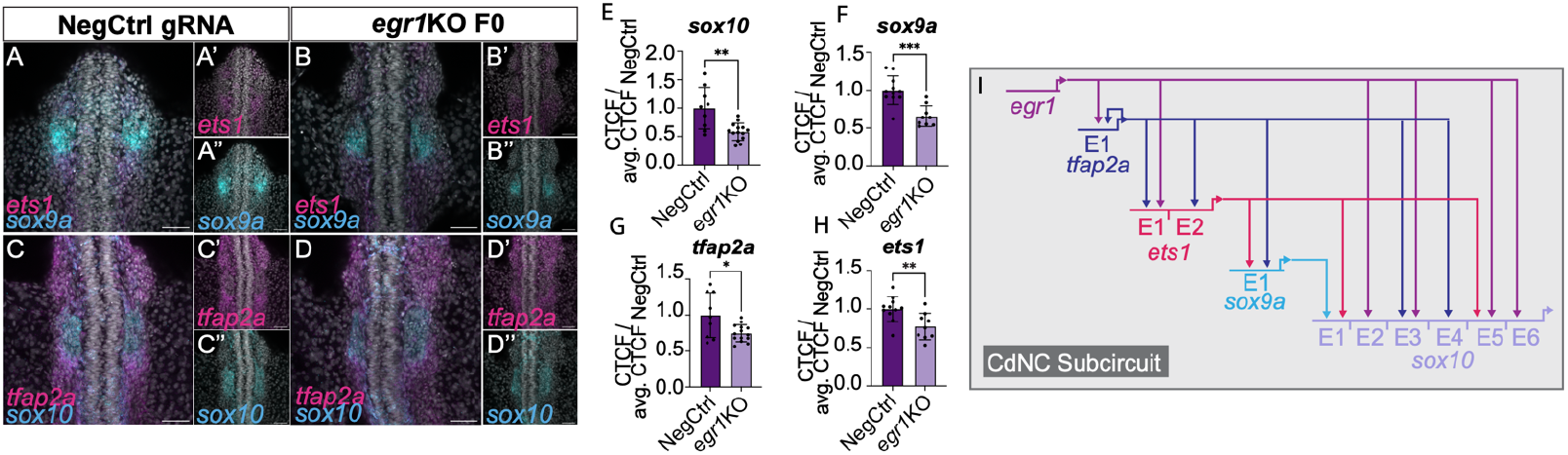
Loss of *egr1* leads to a downregulation of CdNC genes and suggests an important role as core CdNC subcircuit transcription factor. HCR staining for *ets1* and *sox9a* in 16ss zebrafish embryos injected with negative control gRNA (n=11) **(A-A”)** and *egr1* gRNA (n=9) **(B-B”)**. HCR staining for *tfap2a* and *sox10* in 16ss zebrafish embryos injected with negative control gRNA (n=9) **(C-C’’)** and *egr1* gRNA (n=14) **(D-D”). (E-H)** Quantification of CTCF in negative control and *egr1* gRNA injected embryos (unpaired student’s t test p < 0.05). **(I)** Migratory CdNC subcircuit displaying proposed interactions between shown transcription factors and downstream enhancer regions. All scale bars = 50 um.

Given the abundance of Egr1 motifs in CdNC-specific accessible chromatin regions and its predicted binding sites in many of our putative CdNC enhancers, we sought to test its regulatory role by generating *egr1* knockout embryos using CRISPR/Cas9-mediated mutagenesis. *egr1* and scrambled gRNA control F0 crispants were fixed at 16ss and expression of CdNC genes was analyzed via HCR (Fig. 3A-D’’). Quantification of corrected total cell fluorescence (CTCF) in both pre- and post-otic migratory CdNC streams revealed significant downregulation of *ets1* (n=9), *sox9a* (n=9), *tfap2a* (n=14), and *sox10* (n=14) transcripts in *egr1* crispant embryos compared to controls (unpaired t-test, p<0.05) (Fig. 3E-H, Table S4). These results suggest that *egr1* operates as a key transcriptional regulator of migratory CdNC gene expression. Thus, coupling these results with our putative enhancers and transcription factor motif analysis, we generated a gene regulatory subcircuit characterizing the migratory CdNC (Fig. 3I).

### Migratory CdNC genes are expressed during adult cardiac regeneration

*Sox10* is an excellent marker for the developing NC that persists in some lineages. Given that both *sox10* and *tfap2a* have been shown to be upregulated during cardiac regeneration, we next investigated whether other components of our CdNC gene regulatory subcircuit also play a role in cardiac regeneration^2,17^. To test this, we employed weighted gene coexpression network analysis (WGCNA) on bulk RNA-seq data to identify correlated patterns of gene expression between the migratory CdNC and *sox10+* cells in adult regenerating hearts^2,36^. Included in this WGCNA are previously published bulk RNA-seq datasets collected from *foxd3*+ cells in 75% epiboly-stage and 6ss embryos, as well as datasets from migratory 16ss CdNC cells (this paper) and 21 days post amputation (dpa) *sox10*-expressing cells^2,37^. Of the 17 gene expression modules identified, 2 were positively correlated between the migratory CdNC and 21dpa *sox10*+ cells (Fig. S4A). Several genes that we had identified as markers of the migratory CdNC, such as *sox10, egr1, fli1*, and *cxcr4b*, were in one shared coexpression module (“Black” Module, 7064 genes) (Fig. S4B).

To validate our WGCNA findings and further investigate CdNC gene expression during regeneration, we performed HCR for genes in our regulatory subcircuit on wild-type adult ventricles that were either resected or sham-injured. Strikingly, at 7dpa, we found that *egr1* (n=15/15), *ets1* (n=11/11), and *sox9a* (n=12/12) were all expressed in the regenerating tissue, as identified by the reduction of *myl7* expression (Fig. 4A-F’). Transcripts for these genes can be observed in late regeneration as well (21dpa), albeit at lower levels (Fig. S5). Further, we show that *egr1, sox9a*, and *ets1* are also coexpressed in a subset of the regenerating tissue (Fig. 4G-G’’’) (n= 8/8). We next wondered whether *sox10*-expressing cells coexpress these other CdNC genes as they do during developmental migration. Using HCR, we observed colocalization of *egr1* and *sox10* transcripts in a subset of cells after injury (n= 10/11) (Fig. 4H-H’’’). The colocalization of these CdNC genes after injury therefore suggests that they could potentially operate in a regulatory subcircuit as they do during CdNC migration.

**Figure 4.**
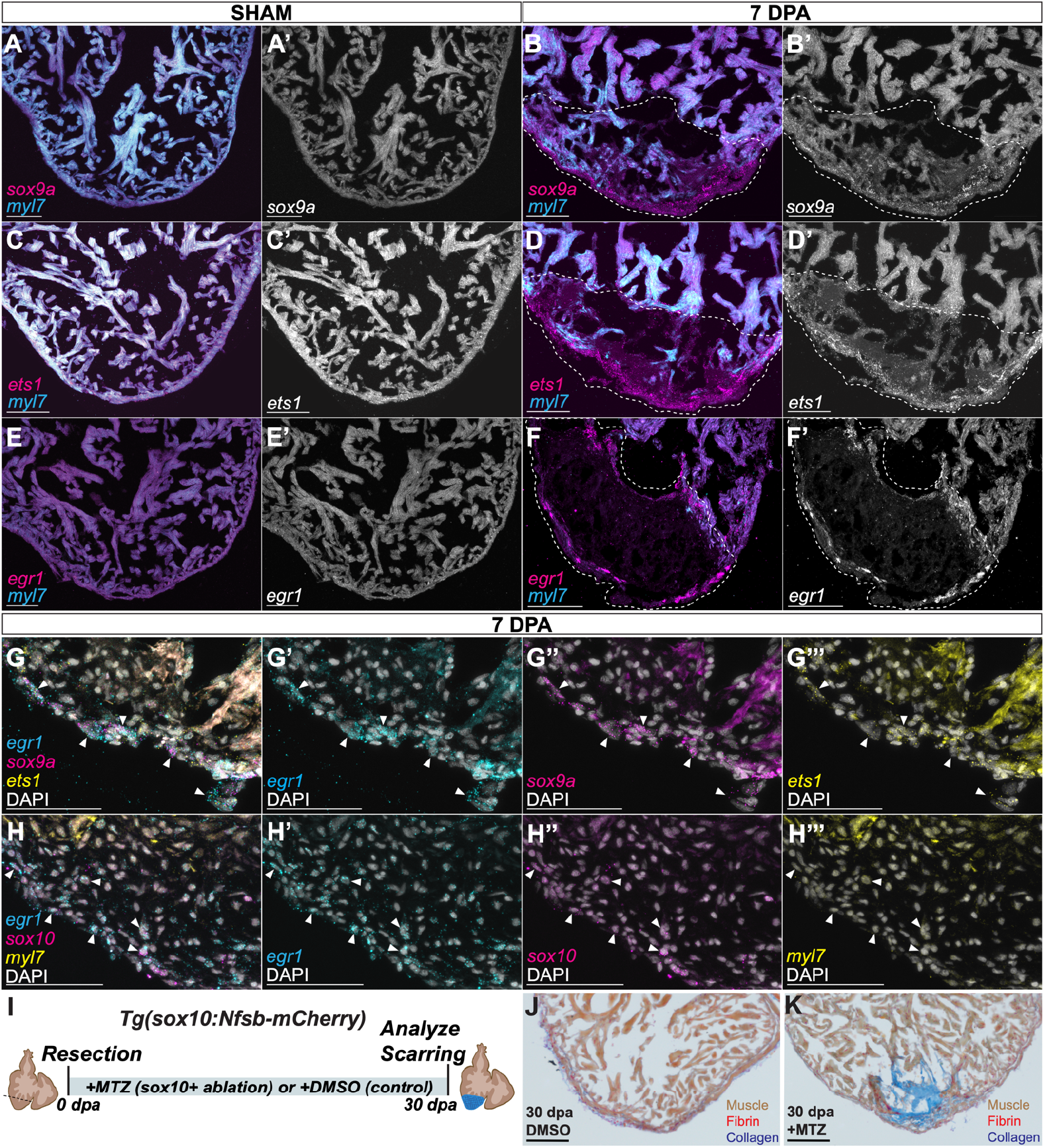
A developmental CdNC subcircuit is expressed during adult cardiac regeneration. HCR staining of sham and injured 7dpa ABWT hearts, respectively, for *myl7* and **(A, B)** *sox9a* (n=12/12), **(C, D)** *ets1* (n=11/11), and **(E, F)** *egr1* (n=15/15). **(A’-F’)** Grayscale of respective *sox9a, ets1*, and *egr1* stainings. Scale bars = 100 um. **(G)** HCR co-staining on ABWT 7dpa hearts for *sox9a, egr1, ets1*, and DAPI (n=8/8). Arrows indicate some of the cells positive for all three probes. Single channel images, with DAPI, of, *sox9a, egr1*, and *ets1* are shown respectively in **(G’-G’’’)** Scale bars = 50 um. **(H)** HCR co-staining on ABWT 7dpa hearts for *sox10, egr1, myl7*, and DAPI (n=10/11). Arrows indicate a subset of the cells double-positive for *sox10* and *egr1* probes. Single channel images, with DAPI, of *egr1, sox10*, and *myl7* are shown respectively in **(H-H’’). (I)** Diagram of the *sox10*+ ablation scheme. Tg(*sox10:Gal4;UAS-Nfsb*) injured fish are maintained in MTZ for the duration of the regeneration time course and hearts are harvested at 30 dpa. **(J)** Injured hearts maintained in DMSO to 30 dpa retain little scarring (n=3/3), whereas **(K)** injured hearts maintained in MTZ, ablating *sox10*+ cells, maintain a scar to 30 dpa (n=5/5). Scale bars = 100 um.

### Genetic ablation of *sox10*-expressing cells impairs adult cardiac regeneration

Finally, we asked if *sox10*-expressing cells were essential for cardiac regeneration in adult zebrafish. We and others have demonstrated that both *sox10*-derived and *sox10*-expressing cells contribute to adult cardiac regeneration in the zebrafish^2,17^. To definitively determine whether cells expressing *sox10*, a central node in our CdNC subcircuit, are necessary for cardiac regeneration, we utilized the genetic ablation line *Tg(sox10:Nfsb-mCherry)* that expresses the bacterial nitroreductase enzyme under control of the *sox10* promoter^11^. Upon resection of 20% of the ventricle, adult *Tg(sox10:Nfsb-mCherry)* fish were maintained in either DMSO or metronidazole (MTZ) throughout the regeneration time course to induce cell death of *sox10*-expressing cells both at the time of injury and throughout regeneration^38^ (Fig. 4I). At 30 dpa, hearts were collected and ventricles assayed for scarring in both MTZ and DMSO-control conditions. Histological assays for scarring were done using acid fuchsin orange G (AFOG), a sensitive stain for fibrin/collagen. By 30 dpa, control hearts regenerate with minimal scarring, indicative of successful regeneration (n=3/3), whereas hearts from the *sox10+-* ablated condition harbored large areas of scarring (n=5/5), suggesting a lack of regeneration (Fig. 4J-K). These findings, using the ventricular resection model, definitively show that cells expressing *sox10* at the time of injury and throughout the regeneration time course are necessary for proper cardiac regeneration. These data complement and are consistent with findings where ablation of *sox10+* cells prior to cryoinjury impaired regeneration^17^. Together, these data support the hypothesis that cells expressing CdNC genes play a critical role in zebrafish cardiac regeneration.

## DISCUSSION

While much progress has been made toward understanding the migratory paths and downstream derivatives of the CdNC, little is known about the regulatory networks that control this unique axial population. In this study, we use a systems-level genomic analysis combined with *in vivo* testing to reveal a gene regulatory subcircuit characteristic of migratory CdNC cells. We identify a novel role for *egr1* in promoting expression of canonical CdNC transcription factors. Importantly, using the zebrafish as a model, we demonstrate that genes within this embryonic gene subcircuit are reactivated after injury in the adult heart, highlighting a potential link between CdNC regulatory networks and adult regeneration. However, further dissection of regulatory linkages between CdNC genes in the injured adult heart is needed to determine if the subcircuit in its entirety is reactivated in response to damage.

*egr1* (early growth response 1), previously known as *krox24*, encodes a highly conserved zinc-finger transcription factor expressed in a variety of contexts in both embryonic and adult tissues^39–42^. Despite previous literature demonstrating its necessity in other tissues, very little has been known about the role of *egr1* in either NC or cardiovascular development. Our data demonstrate that knockout of *egr1* leads to the downregulation of critical CdNC genes, including *tfap2a, sox9a, ets1*, and *sox10*, at migratory stages. These data further support our putative GRN subcircuit, but future experiments are required to uncover direct transcription factor binding to our CdNC enhancers. Unlike *egr1*, perturbation of other genes in the subcircuit have documented cardiac phenotypes. The loss of *Ets1* in frogs and mice leads to dysregulated CdNC migration and differentiation, resulting in various cardiac defects^43–45^. *Sox9* expression in murine CdNC cells is critical for the development of the endocardial cushions in the distal part of the outflow tract^46^.

Importantly, *egr1* has been shown to play a role in a variety of disease and injury models through regulating fibrosis and wound healing, but the impact is context-specific. For instance, overexpression of *Egr1* in mice enhances wound healing of incisional wounds, while studies in rat and pig have shown that *Egr1* inhibition reduces the pathological effects of myocardial infarction^47–50^. More recently, one group demonstrated that *Egr1* KO in both adult and neonatal hearts impairs regenerative senescence and heart regeneration in mice^51^. In more extreme cases of whole body regenerative organisms, acoel flatworms with *egr* knockdown fail to regenerate, suggesting a critical role for *egr* in regeneration throughout the animal kingdom^52^. Thus, the role of *egr1* in regeneration is complex and likely dependent on the role that fibrosis plays in different injury paradigms.

Interestingly, though *egr1* was assigned to the CdNC-like module in our WGCNA analysis (correlation = 0.898, Fig. S4B), it was also highly correlated with the other shared module between our development and regeneration datasets (correlation = 0.732), which contains key fibroblast genes like *postnb* (Fig. S4C). This observation raises the intriguing possibility that *egr1* is expressed in a subset of cardiac fibroblasts. There is evidence for CdNC-derived fibroblasts in mice, introducing interesting questions regarding the role of *egr1* in CdNC-derived cell types^10^. While no studies have highlighted a role for *egr1* in zebrafish cardiac regeneration, a published gene regulatory analysis of available transcriptomic data from zebrafish heart regeneration predicts *egr1* as a top “master regulator” of regeneration^53^.

Both this and previous work demonstrate that *sox10*-expressing cells play an essential role in zebrafish cardiac regeneration^17^. Furthermore, we show that the CdNC regulatory subcircuit comprising *egr1, sox9a*, and *ets1* is expressed in the regenerating cardiac tissue and that a subset of *egr1*+ cells are also *sox10*+. An open remaining question concerns the role this regulatory subcircuit plays during regeneration, and whether other CdNC genes are also expressed. The reactivation of this CdNC subcircuit in the regenerating heart suggests that CdNC-derived cells may be reacquiring a developmental-like state during regeneration. Support for this hypothesis comes from a recent study of the zebrafish jaw where *sox10* and *sox9a* are re-expressed during repair of craniofacial cartilage. In this system, similar to the cardiac regeneration model, when *sox10*-expressing cells are ablated, regeneration fails^54^.

A plausible alternative to reactivation of CdNC genes being specific to CdNC-derived cells during regeneration is that cells of diverse developmental origins are expressing this CdNC subcircuit during regeneration in order to adopt a CdNC-like phenotype to facilitate repair. NC cells are inherently migratory and multipotent, and cardiomyocytes must undergo significant gene regulatory changes in order to de-differentiate, proliferate, and contribute to lost or damaged myocardial tissue. Further studies involving CdNC-specific conditional gene knockouts are essential next steps to fully understand the relationship between the CdNC, the CdNC GRN, and cardiac regeneration.

Our study provides evidence for an essential role of cells expressing migratory CdNC genes during cardiac regeneration. Given that the failure of these cells to proliferate adversely affects the repair process in zebrafish, we hypothesize that stimulating these cells to reactivate an embryonic program in amniotes, including humans, may open novel avenues to repair damaged heart tissue. Thus, uncovering the GRN that regulates CdNC contributions to the cardiovascular system may facilitate identification of reprogramming therapies aimed at imbuing the human heart with regenerative capacity.

## ACKNOWLEDGEMENTS

We would like to thank Fish Facility staff at Caltech (David Mayorga and Ryan Fraser) and UC Berkeley (Tyler Mentley, Frances Campbell, and Lindsey Arenson) for zebrafish husbandry and care. We would also like to thank Diana Perez and Rochelle Diamond at the Caltech Flow Cytometry Cell Sorting Facility for help with FACS assistance. We would also like to acknowledge the Caltech Millard and Muriel Jacobs Genetics and Genomics Laboratory, in particular, Igor Antoshechkin and Vijaya Kumar for sequencing of our ATAC-seq and RNA-seq libraries. We would also like to thank Dr. Michael Piacentino and Dr. Ezgi Kunttas for dissection assistance for the TNC RNA-seq samples and Dr. Piacentino for consulting on image quantification. We would like to thank Eli Grossman for technical assistance in early gRNA design and synthesis. We would also like to thank Isaac Hilton-VanOsdall for his support in troubleshooting computational pipelines. Thank you to Dr. Iswar Hariharan and Dr. Lara Busby for providing us with feedback on the manuscript.

## AUTHOR CONTRIBUTIONS

The project and experimental design was conceived by M.L.M. and M.E.B. scRNA-seq, bulk RNA-seq, bulk ATAC-seq collection was performed by M.L.M. Analysis of sequencing data was performed by M.L.M., R.M.D-R., and J.K.M. HCR data collection, analysis, quantification, and interpretation was performed by R.M.D-R., A.K.H., L.W.L., M.L.M. Manuscript was written by M.L.M., M.E.B., R.M.D-R., A.K.H., L.W.L. with editing from J.K.M.

## DATA AVAILABILITY

The raw and processed sequencing data has been deposited at NCBI (Bioproject Accession Number: PRJNA1185249). Unprocessed czi files for all confocal images included in this manuscript can be found on Figshare (https://figshare.com/projects/DhillonRichardsonHaugan_PNAS/227391). Code for data processing and analysis can be found on Github (https://github.com/Martik-Lab/DhillonRichardsonHaugan_PNAS2024).

## METHODS

### Zebrafish

Zebrafish experiments in this study used the wild-type AB strain and transgenic lines *Tg(−4.9sox10:GAL4-UAS-cre;UAS:NfsB-mCherry;myl7:nucGFP)*^11^ and *Tg*(*-4.9sox10:eGFP)*^20^. All adult regeneration experiments were done with fish or random sex and older than 3 months. Adult fish were maintained in a 14/10 hour light/dark cycle, maintained at 28°C, and fed twice daily. All experiments were approved and complied with the California Institute of Technology and UC Berkeley Institutional Animal Care and Use Committee (IACUC) protocol 1764 and AUP-2021-03-14107-1, respectively.

### Bulk RNA-seq and ATAC-seq of Migratory NC Cells

To collect migratory CdNC samples for bulk RNA-seq, *Tg(−4.9sox10:GAL4-UAS-cre;UAS:NfsB-mCherry;myl7:nucGFP)*^11^ 16ss embryos were dissected, isolating the region located between the midbrain-hindbrain boundary to somite 6. To isolate the migratory TNC, *Tg*(*-4.9sox10:eGFP)*^20^ embryos were dissected at 24 hpf from somite 7 to somite 16. To collect cells for bulk RNA-seq, tissues were dissociated in Accumax (Stem Cell Technologies cat: 07921) at 30°C until a single cell suspension was reached. Cells were subsequently sorted on a BD FACSAria™ Fusion Flow Cytometer. For the migratory CdNC samples, mCherry+ cells were collected as the CdNC population, and mCherry-cells were collected for the non-NC control population (3 replicates for each condition). For the migratory TNC samples, GFP+ cells were collected, with GFP-cells as the non-NC control (3 replicates for each condition). cDNA was prepared from each sample using the SMART-Seq® v4 Ultra® Low Input RNA Kit for Sequencing (Takara cat: 634894) according to the manufacturer’s protocol. Standard Illumina protocols were used to construct each sequencing library and an Illumina HiSeq2500 sequencer at the Millard and Muriel Jacobs Genetics and Genomics Laboratory (California Institute of Technology, Pasadena, CA) for 50 million, single-ended reads. For bulk ATAC-seq, CdNC populations were isolated from the same transgenic line and dissections as for bulk RNA-seq. Samples were then dissociated, sorted, libraries prepped, and sequenced according to our previously published protocol^55^. Briefly, tissue samples were dissociated on ice using a cold protease cell dissociation buffer until a single cell suspension was achieved. Positive and negative populations of migratory CdNC and TNC were collected using the same fluorescent cell sorting scheme as the bulk RNA-seq. Cells were then lysed with a cold lysis buffer, regions of open chromatin fragmented and tagged using a Tn5 transposase, and sequencing libraries were constructed using barcoded primers. Bulk ATAC-seq libraries were sequenced on an Illumina HiSeq2500 sequencer at the Millard and Muriel Jacobs Genetics and Genomics Laboratory (California Institute of Technology, Pasadena, CA) for 50 million, paired-end reads.

### Bulk ATAC-seq processing and analysis

To analyze the bulk ATAC-seq datasets, FastQC was used to perform initial quality checks after sequencing. Sequencing adapters were then trimmed using Cutadapt v2.8^56^. Trimmed, paired-end reads were mapped to the zebrafish genome (GRCz11) using Bowtie2^57^. Samtools (view -s) was used to downsample each replicate to approximately 95 million reads^58^. PCR duplicates and mitochondrial reads were filtered out and peaks were called with Genrich. bamCoverage was used to generate Bigwig files from downsampled bam files for visualization in IGV (−bs 10, –normalizeUsing RPGC). Peaks differentially accessible between samples were identified using the standard DiffBind workflow^30,31^. The Diffbind output was used as an input to VolcaNoseR to create volcano plots^59^. These differentially accessible regions were manually extended to encompass the entire peak length and genomic coordinates were determined in IGV and downloaded using Ensembl. HOMER^32^ was used to discover *de novo* motifs in the regions of chromatin differentially accessible in the CdNC sample. A background bed file with regions differentially accessible in either the CdNC sample or the negative sample was provided to HOMER. Putative enhancers were then scanned for transcription factor binding motifs with FIMO^34^, using the 2024 JASPAR non-redundant vertebrate motifs dataset^60^. In FIMO, we used the default background model and filtered out any binding sites with a p-value <1E-4. Enhancer box plot diagrams were constructed to relative scale in Adobe Illustrator.

### Analysis of Bulk RNA-seq and WGCNA

To analyze the CdNC and TNC bulk RNA-seq datasets, reads were mapped to the zebrafish genome (GRCz10) using Bowtie2^57^. Transcript counts were calculated using featureCounts (Subread) and differential gene expression analysis was performed using DESeq2^61^. Volcano plots were produced using fold change and p-value measurements from DESeq2 using VolcaNoseR^59^. To identify coexpression modules between development and regeneration, we utilized the R package, WGCNA^36^, on bulk RNA-seq from 21dpa *sox10*+ cells (previously published)^2^, 16ss CdNC cells, 6ss embryonic cells, and epiboly stage embryonic cells (previously published)^37^. A soft threshold power of 20, a minimum module size of 100, and a module merging threshold of 0.25 were used for this analysis.

### scRNA-seq Analysis of Migratory CdNC Cells

To collect scRNA-seq data from migratory CdNC cells, *Tg(−4.9sox10:GAL4-UAS-cre;UAS:NfsB-mCherry;myl7:nucGFP)*^11^ 16ss embryos were dissected, dissociated, and FAC-sorted. Single cell libraries were prepared using the Chromium Single Cell 3’ v3.1 reagent kit (10X Genomics, PN-1000075). 3,828 cells were sequenced, creating an average read depth of 80,000 reads per cell. Reads were mapped to the zebrafish genome (GRCz11) and low quality cells were excluded, using 10x Genomics Cell Ranger v7.20. Scanpy was used for further quality control, filtering, clustering, and downstream analysis^27^. Cells were filtered by n_genes_by_counts, total_counts, percent_counts_mitochrondrialgenes, and percent_counts_ribosomalgenes. Remaining mitochondrial, ribosomal, and cell cycle genes were regressed out prior to downstream analysis. Principal component analysis was performed and a nearest neighbor graph was computed (scanpy.pp.neighbors). This graph was embedded in two dimensions using UMAP (sc.tl.umap). Leiden clustering was performed (sc.tl.leiden) and visualized (sc.pl.umap). Marker genes were calculated based on the Wilcoxon rank sum (sc.tl.rank_genes_groups) and plotted (sc.pl.rank_genes_groups). Clusters were annotated based on the expression of marker genes. velocyto, and R package, was used to create loom files for velocity analysis and scVelo was used for further downstream RNA velocity analysis^28,62^.

### Hybridization Chain Reaction of Embryonic Zebrafish

Zebrafish embryos were staged and fixed in 4% PFA in PBS for 2 hours at room temperature or overnight at 4°C, followed by PBST washes and dehydration in 100% ethanol. Hybridization chain reaction was performed using molecular instruments whole-mount zebrafish embryos and larvae protocol with minor modifications. Briefly, embryos were rehydrated and manually dechorinated in graded EtOH/PBSTw washes and hybridized with probes with concentrations varying from 2 -8 pmols depending on the probe. The following day, probes were thoroughly washed with probe wash buffer and 5X SSCT and amplified with 30pmol of individually snap-cooled hairpins overnight. Lastly, the embryos were thoroughly washed with 5x SSCT and incubated in 1:1000 DAPI for 30 minutes. Following HCR, embryos were mounted in 0.40um canyon molds and immersed in 5x SSCT. Images were taken with a 20x dipping lens on a Zeiss 780 LSM upright confocal microscope, and image analysis was performed using Fiji. All images are displayed as maximum intensity projections of all z-stacks. All probes, hairpins, and buffers were purchased through Molecular Instruments. The following probes were used in zebrafish embryos: B4 or B5 *sox10*, B3 *cxcr4b*, B3 *prdm1a*, B2 *mafba*, B7 *foxc1b*, B7 *tfap2a*, B9 *twist1a*, B4 *twist1b*, B2 *nrp2b*, B8 *fli1*, B1 *ets1*, B6 *sox9a*, B10 *tgif1*.

### *egr1* gRNA Design and Injection

To generate high-efficiency edits in F0 zebrafish, we followed a previously published protocol with slight modifications^63^. Synthetic guide RNAs (gRNA) were designed utilizing the Alt-R CRISPR editing system created by Integrated DNA Technologies (IDT). We took the top three gRNAs for the zebrafish *egr1* gene selected from IDTs database of predesigned crRNAs and ran them through inDelphi, a machine learning algorithm which predicts gRNA efficiency and frameshift frequency^64^. From this, we found one crRNA upstream the DNA-binding domains of *egr1* that had a frameshift frequency of 83.7% with the most likely edits being a 1-bp insertion and an 8-bp deletion. Already established scrambled crRNAs from IDT were purchased as a negative control for HCR quantification. Sequences for the scrambled and *egr* crRNAs can be found in supplementary table 3. To generate a single gRNA, 1uL crRNA, 1uL tracrRNA, and 1.51uL Duplex Buffer (supplied by IDT) were combined and annealed at 95°C for 5 minutes and then cooled on ice, resulting in a final gRNA concentration of 28.5uM. Cas9 protein (wild-type S. pyogenes Cas9 protein with a double NLS tag (SV40-derived) at the C-terminus purchased through UC Berkeley QB3 Macrolab or IDTs Alt-R S.p. Cas9 Nuclease V3 cat: 1081058) was diluted to 30uM in dilution buffer, consisting of 20 mM Tris-HCl, 600 mM KCl and 20% glycerol. Given that preassembled ribonucleoproteins (RNPs) have shown higher efficiency in generating edits in zebrafish, we combined equal volumes of gRNA and diluted Cas9 protein, incubated them at 37°C for 5 minutes, and kept on ice for injection or kept at -20°C for long-term storage^65^. For injections, about 1nL of RNP solution was injected into the yolks of single cell stage of zebrafish embryos prior to cell inflation. Injected embryos developed until 16ss, fixed in 4% PFA for 2 hours at RT or overnight at 4°C, dehydrated in 100% EtOH, and stored at -20°C until HCR.

### *egr1* Crispant Genotyping and Image Quantification

Following HCR, crispant embryos were mounted and imaged as described in the zebrafish embryo HCR section. To compare expression between crispant and negative control embryos, laser intensity and gain remained constant between all embryos during imaging. Initial settings were calibrated by analyzing the expression intensity of each channel in 3 negative control embryos. Following imaging, we individually genotyped each embryo. To isolate genomic DNA (gDNA), we followed a standard protocol with modifications^66^. Whole embryos were placed into labeled PCR strip tubes containing 25uL lysis buffer (for 50mL, combine 500uL 1M Tris-HCl, pH=8.3, 2.5mL 1M KCl, 75uL 1M MgCl2, 1.5mL 10% Tween-20, 1.5mL 10% NP40, and ddH2O up to 50mL) and 2.5uL of 20mg/mL proteinase K (Thermo scientific cat: EO0491). Tubes were incubated at 55°C for 180 minutes and 94°C for 20 minutes. 1uL of gDNA was used with 0.5uM of forward (5’-TGGACCGGAGATGAGTAGCA-3’) and reverse (5’-CCTCTGTTCAGCCTGGTGAG-3’) primers. This region was amplified using Q5 HotStart 2x High Fidelity Master mix (cat: M0494X), and the samples were submitted for sanger sequencing. We compared the amplicons from negative control and *egr1* crispant embryos using TIDE, a method for identifying the editing frequency and mutation class^67^. Using these results, we only quantified the expression of *egr1* crispant embryos which contained >40% editing efficiency for insertions or deletions that would result in a frameshift mutation. Additionally, we noticed many of our embryos contained high proportions of 1-bp insertions and 8-bp deletions, suggesting that InDelphi is an accurate tool for selecting promising gRNAs (TableS 3). Image quantification of genotyped embryos was performed in Fiji on maximum intensity projections of all z-stacks. We followed a previously published quantification scheme with some modifications^68^. The migratory CdNC regions of interest (ROIs), encompassing both the pre and post otic streams of neural crest cells were drawn based on *sox10* expression. Additional regions, to be excluded from ROI calculations, encompassing the otic vesicles were drawn based on embryonic morphology. Calculated total cell fluorescence (CTCF) measurements were first determined for the entire migratory CdNC ROI (including the otic vesicles) as followed:

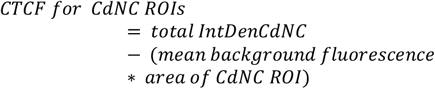

Then, CTCF measurements were determined for the otic vesicles as followed:

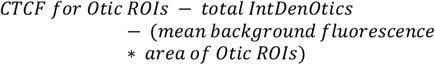

Finally, we subtracted the CTCF values of the otic vesicles from the CTCF value of the entire CdNC ROI to calculate the fluorescence intensity value for only migratory CdNC cells (excluding signal from the otic vesicles). Unpaired student t-test between control and CRISPR-treated experimental embryos was used to determine statistical significance between the two populations (p<0.05).

### Ablation of *sox10*+ Cells During Cardiac Regeneration

To inducibly ablate *sox10*-expressing cells, we utilized the nitroreductase genetic cell ablation system^69^ with the line *Tg(−4.9sox10:GAL4-UAS-cre;UAS:NfsB-mCherry;myl7:nucGFP)*^11^ (referred to as *Tg(sox10:Nfsb-mCherry)* in the text). After cardiac resection, as described in the methods, adult zebrafish were maintained in either 5 mM Metronidazole (Sigma Aldrich cat: M1547) or DMSO for the duration of the regeneration time course. Water with fresh drugs was refreshed at least 1x/day. After 30 days, hearts were harvested as described in the methods and processed for AFOG staining.

### Acid Fuchsin Orange G (AFOG) Staining

After ablation and tissue fixation, *Tg(−4.9sox10:GAL4-UAS-cre;UAS:NfsB-mCherry;myl7:nucGFP)* hearts were embedded in paraffin and sectioned at 10 uM thickness on a Zeiss microtome. AFOG staining was performed using a modified Masson’s trichrome procedure^38^ (Sigma Diagnostics Procedure No. HT15). Briefly, slides are hydrated in water for 10 minutes, then incubated in Bouin’s Solution at 60°C for 2 hours, followed by an additional hour at room temperature. Slides are rinsed in water for 30 minutes at room temperature, rinsed in 1% Phosphomolybdic acid for 5 minutes, followed by a 5 minute water rinse. Slides are incubated in AFOG staining solution for 5 minutes, rinsed with water for 5 minutes, and then dehydrated. Lastly, sections are cleared with Xylene and mounted with Cytoseal 280 (Thermo Fisher cat: 8311-4).

### Zebrafish Cardiac Injury and Tissue Collection

Adult zebrafish heart resection was conducted on ABWT and *Tg(sox10:NfsB-mCherry)* ventricles according to published protocols^38^. Adult fish were anesthetized in 200 mg/mL MS-222 (Millipore Sigma cat: A5040) and up to 30% of each ventricle was amputated. Resected and sham operated hearts were collected at 7 and 21 days post amputation (dpa) at which time the fish were euthanized. Hearts were extracted, placed in heparin buffer on ice for 5-15 minutes, followed by perfusion buffer for 1-4 hours on ice. Recipes for both heparin and perfusion buffers are from Sander et al. 2013, with the penicillin-streptomycin omitted^70^. While in the perfusion buffer, hearts were carefully mechanically perfused using forceps. Once perfusion is complete, hearts are delicately punctured with 30 gauge needles to allow paraformaldehyde to penetrate the entire heart.

Hearts were fixed in 4% PFA in PBS overnight at 4°C while gently shaking. For cryosectioning, hearts were then washed in PBS and put through 5% and 15% sucrose steps (until hearts equilibrate) and embedded in OCT on dry ice for cryosectioning. For whole-mount HCR, hearts were briefly washed in PBS and dehydrated in 100% EtOH. Following whole mount HCR, hearts were embedded in plastic using the JB-4 plastic embedding kit and its published protocol (Sigma-Aldrich cat: EM0100). Embedded hearts were sectioned at 5-8um thickness on a microtome and mounted in Fluromount-G® (Southern Biotech cat: 0100-01) for confocal microscopy.

### Hybridization Chain Reaction on Adult Zebrafish Hearts (Cryosections and whole mount)

Hybridization Chain Reaction (HCR) was performed on cryosectioned zebrafish hearts using the Molecular Instruments HCR™ RNA-FISH ‘fresh/fixed frozen tissue samples’ protocol. The following probes, all designed and manufactured by Molecular Instruments, were used for HCR on adult zebrafish heart cryosections: *myl7* B4, *myl7* B5, *sox9a* B1, *sox9a* B6, *ets1* B1, *egr1* B3. HCR was also performed on whole mount zebrafish hearts. The following probes were used for whole-mount zebrafish hearts: B3 *egr1*, B5 *myl7*, and B4 *sox10*. Heart sections were subsequently imaged with confocal microscopy. For all injured and sham gene expression comparisons, all laser settings and image processing is kept the same.

---

**Fig. S1.**
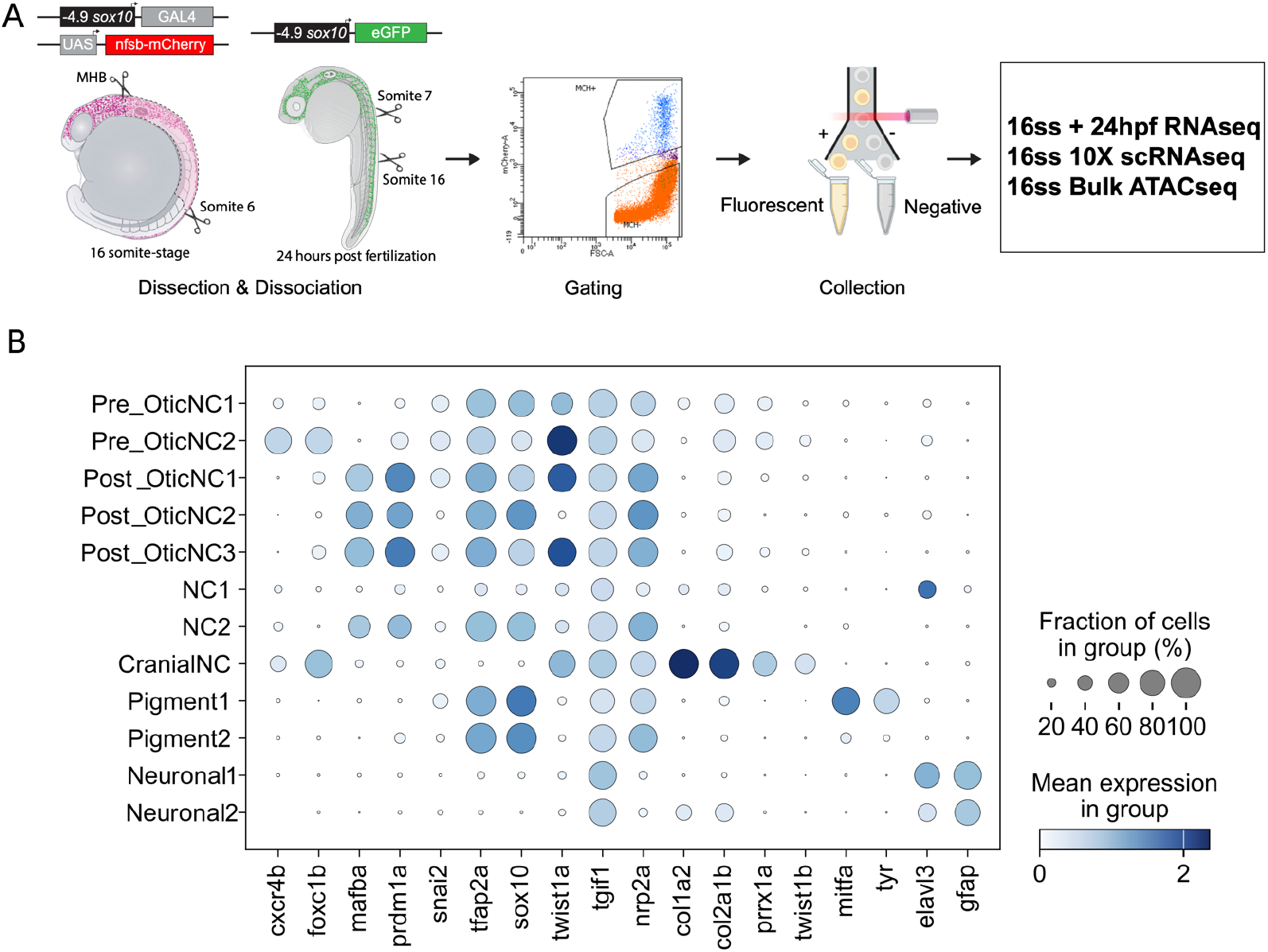
Sequencing dataset collection and scRNA-seq marker genes. **(A)** Schematic of all sequencing datasets collected, beginning with tissue dissections and cell dissociations of embryonic transgenic zebrafish, fluorescence-activated cells sorting (FACS) to isolate pure populations of NC cells, and sequencing (illustrations made with BioRender). **(B)** Dot plot showing the mean expression and proportion of specific genes in each scRNA-seq cluster for cluster identity validation.

**Fig. S2.**
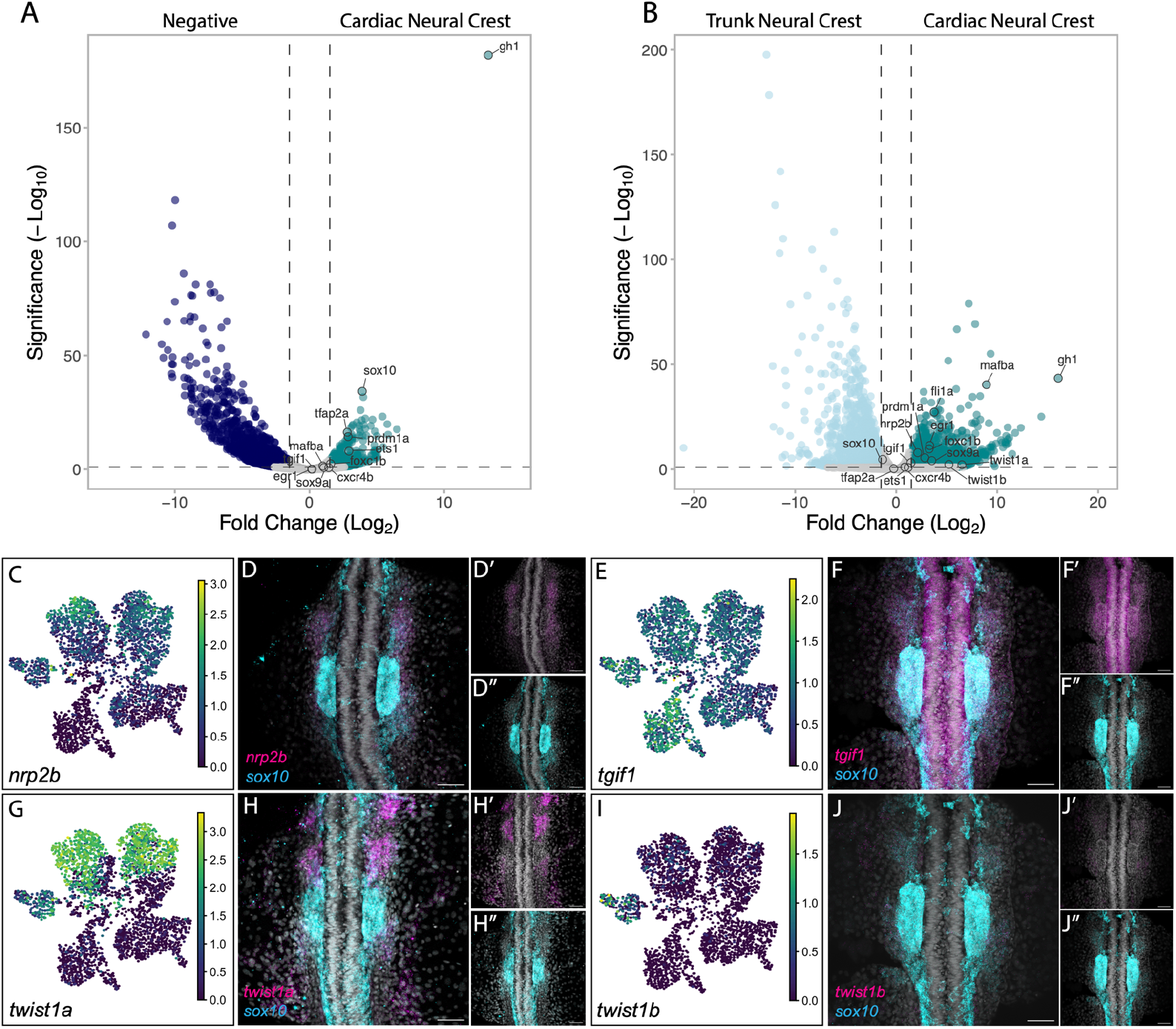
Differentially expressed genes in sorted CdNC cells and additional HCR validations. **(A-B)** Volcano plot of bulk RNA-seq data showing genes differentially expressed in CdNC versus negative/non-NC samples **(A)** CdNC versus TNC samples **(B). (C-J’’)** UMAPs for individual genes and corresponding 20x confocal images of HCR expression patterns of 16ss zebrafish embryos, showing overlap in expression and individual channel insets of *sox10* in cyan and the following genes of interest in magenta: **(C-D’’)** *nrp2b* (n=3), **(E-F’’)** *tgif1* (n=4), **(G-H’’)** *twist1a* (n=8), **(I-J’’)** *twist1b* (n=4).

**Fig. S3.**
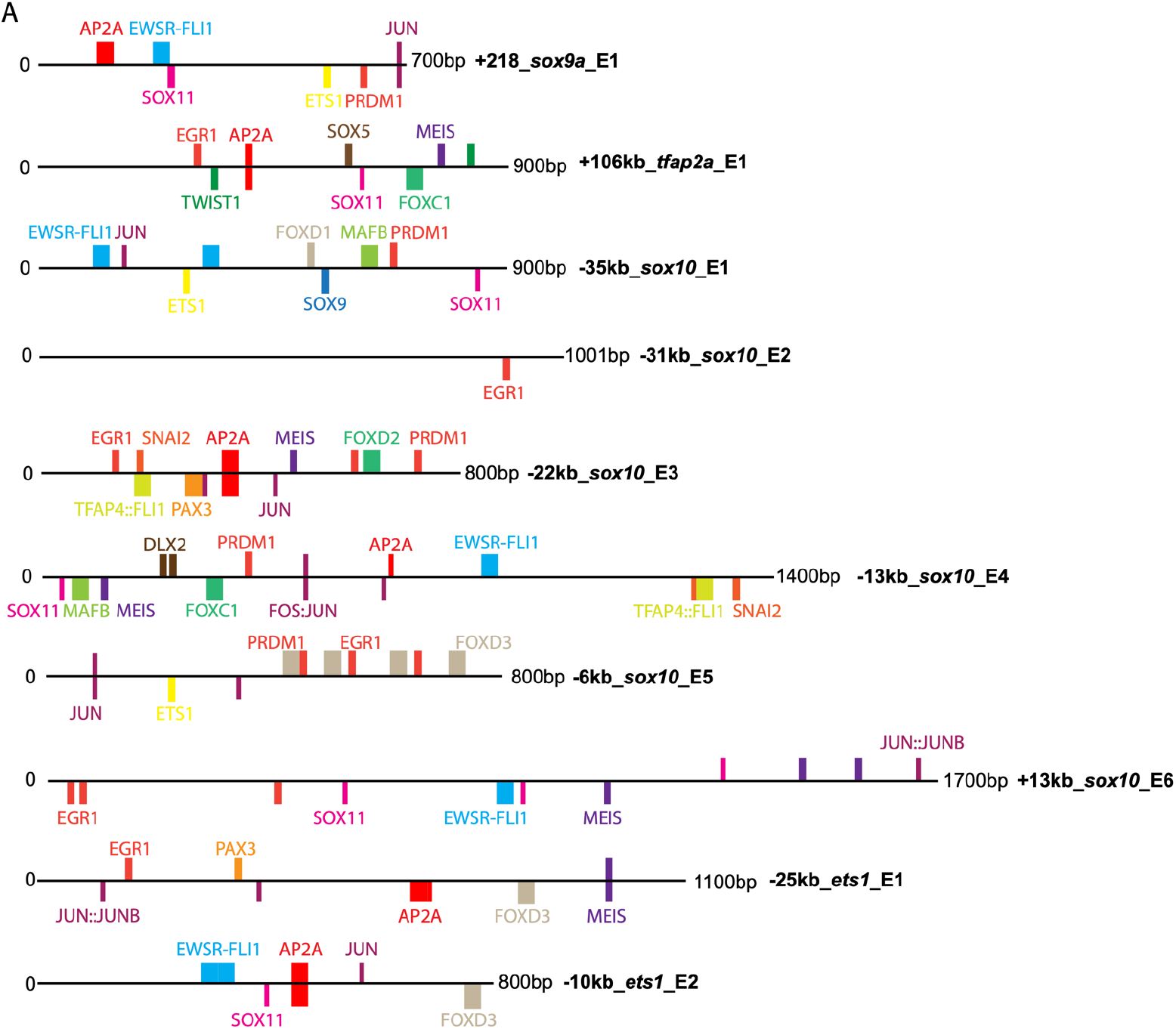
ATAC transcription factor binding site analysis. **(A)** Box plot diagrams of putative enhancers in our CdNC subcircuit (*tfap2a* E1, *ets1* E1 and E2, *sox9a* E1, and *sox10* E1-E6). Relevant NC motifs were annotated based on FIMO output. Transcription factor binding sites less than 10 base pairs are represented by shorter bars and binding sites greater than 10 base pairs are represented by the larger bars.

**Fig. S4.**
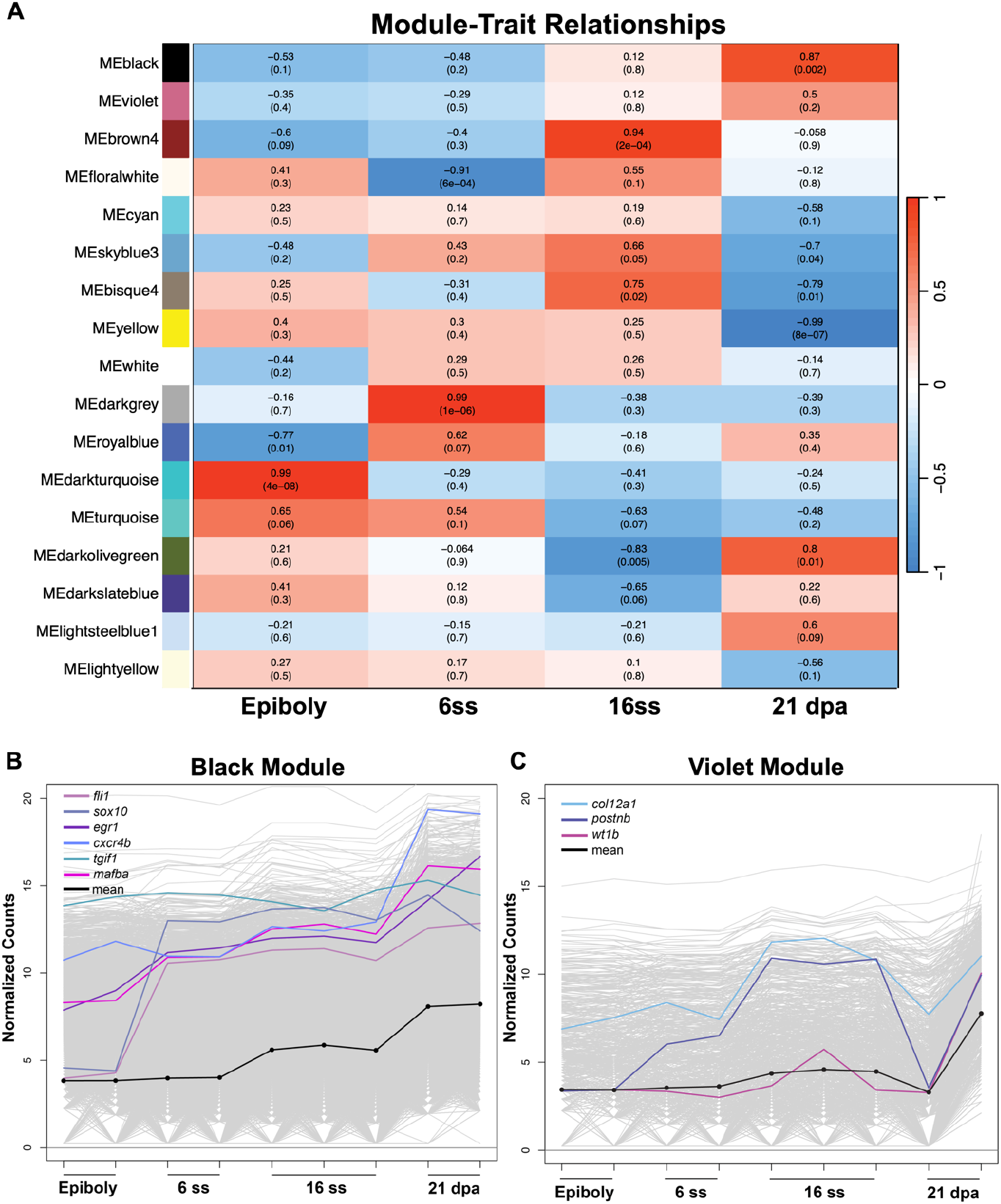
WGCNA Analysis between migratory CdNC and regenerating 21dpa *sox10+* cells. **(A)** Table of WGCNA modules with their correlation and associated p-value (in parentheses) to each bulk RNA-seq time point. **(B)** Line plot of normalized counts for each gene in the WGCNA module shared between migratory CdNC and regeneration (“Black” Module) with key migratory genes highlighted. **(C)** Line plot of normalized counts for each gene in the violet module, with key fibroblast/epicardial genes highlighted.

**Fig. S5.**
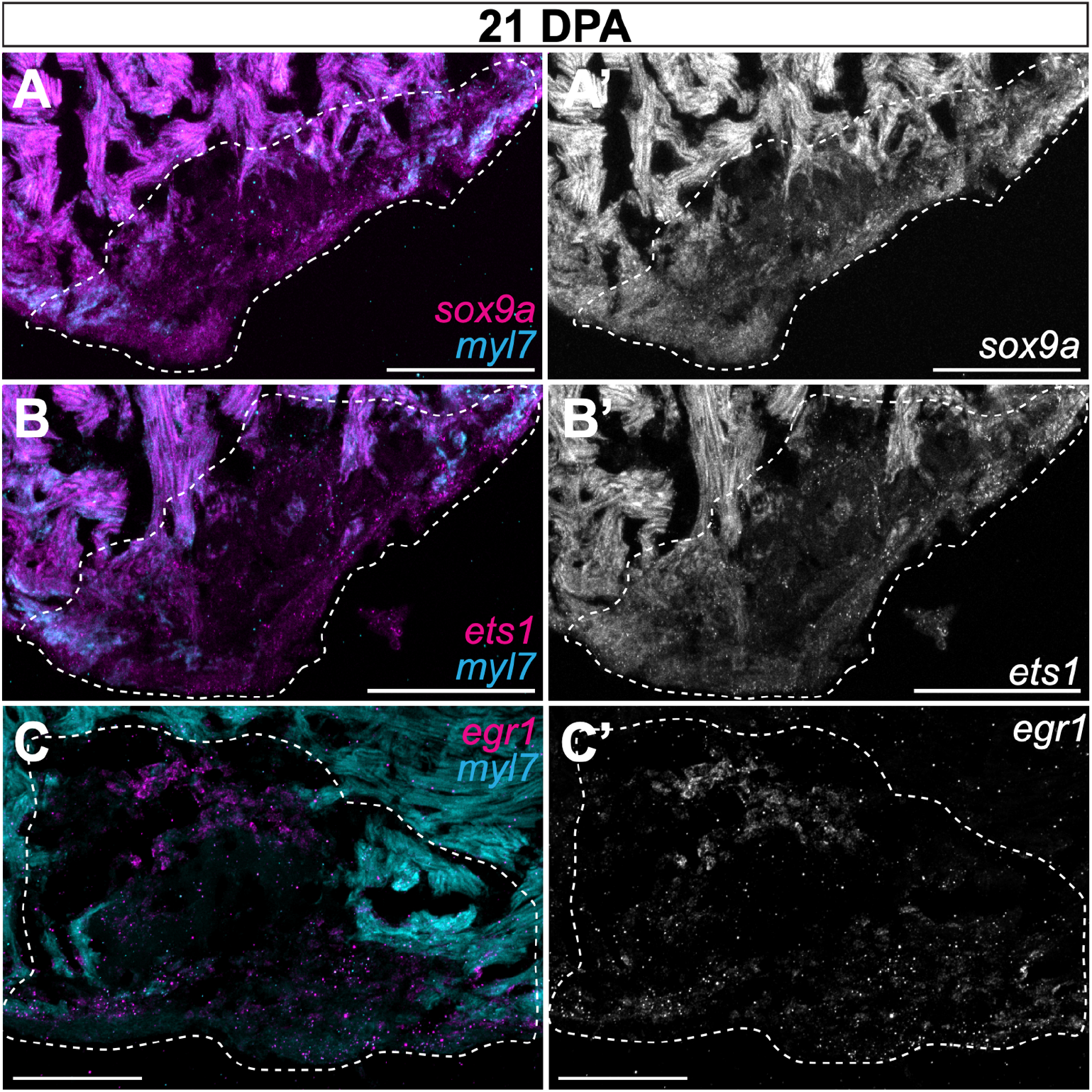
CdNC gene expression at 21dpa in the adult heart. HCR staining of the regenerating adult ventricle at 21dpa with *myl7* and **(A-A’)** *sox9a*, **(B-B’)** ets1, and **(C-C’)** *egr1*. Approximate regenerated area is outlined. Scale bars = 100 um.

**Table S1**: Bulk RNA-seq differentially expressed genes

All differentially expressed genes of bulk RNA-seq data between CdNC versus Non-NC/Negative as well as CdNC versus Trunk NC

**Table S2**: DiffBind output and putative enhancers

Diffbind output for all called differentially accessible regions of open chromatin

**Table S3**: Putative enhancer coordinates and FIMO results

GRCz11 coordinates, DNA sequence, and FIMO output for all putative CdNC enhancers

**Table S4:** gRNA sequences and crispant genotyping

gRNA sequences and genotyping primers for *egr1* and negative control crispant experiments, TIDE output of top three INDELs generated for each embryo and corresponding CTCF values of quantified HCR images.

